# Consistent visuomotor adaptations and generalizations can be achieved through different rotations of robust motor modules

**DOI:** 10.1101/362228

**Authors:** Cristiano De Marchis, Jacopo Di Somma, Magdalena Zych, Silvia Conforto, Giacomo Severini

**Author notes:** Corresponding Author: Giacomo Severini, School of Electrical & Electronics Engineering, University College Dublin, Belfield, Dublin 4, Ireland.

## Abstract

Humans have a remarkable ability to modify their motor commands in response to alterations in the environment. These motor commands are thought to be adaptively tuned by combining different motor primitives, a characteristic that allows for adaptations to be generalized also to relevant untrained scenarios. A complementary theory of motor primitives has shown that natural movements can be described by the combination of relatively invariant sets of muscle synergies. Synergistic structures have been shown to be maintained and tuned during motor adaptations. Here we studied the influence of synergistic organization of movement at the muscular level on the way in which adaptations to a 45° clockwise visuomotor rotation are achieved and generalized. We confirm that adaptation is achieved by tuning a set of robust muscle synergies and we show that the same biomechanical adaptation can be achieved by differentially tuning the same synergies. We demonstrate that this differential tuning influences the presence and extent of generalization and depends on the way in which the targets are experienced relatively to the synergistic structure itself. Our results support the speculation that muscle synergies may be a part of the functional motor primitives that the motor system updates during motor adaptations.

## Introduction

Complex movements can be easily adapted in response to discrepancies between the expected and actual sensorimotor output. This is achieved through the recalibration of the neural representation of the body and the environmental interaction dynamics that are thought to be stored in internal models that characterize both the forward and inverse dynamics of movement^1-6^. Internal models can be adapted as a response to changes in the mapping between movement and its associated feedback or in response to perturbations modifying the dynamic characteristics of the movement in a relevant way^7^.

Several seminal studies have suggested that internal models are constituted by combinations of motor primitives^8-12^, an observation that is corroborated by the fact that adaptation to specific perturbations can be generalized to un-trained compatible contexts^9-11,13-16^. These motor primitives are thought to explain the mapping between state variables such as limb position and movement velocities, into motor commands^12^.

A conceptually correlated theory of motor primitives and movement modularity has been proposed and widely demonstrated in both animal models and humans, that is, the muscle synergies hypothesis^17-20^. Muscle synergies are thought to be the functional building blocks of force production during natural movements. Studies employing stimulation^21-24^ and optogenetics^25,26^ techniques have individuated the presence of such modules in the spinal cord of rodents, frogs and primates. In humans, on top of the fertile literature describing muscle synergies during natural behaviors^27-32^, studies on impaired individuals have demonstrated the solidity of this hypothesis in describing neurological conditions such as stroke^33,34^.

As motor adaptations have been mainly studied using computational models derived from metrics of biomechanical error, there is a surprising small evidence connecting the re-organization of the internal models that happens during adaptation with the supposed functional constituents of movement. Although previous studies have suggested muscle synergies as potential physiological blocks of computationally-derived primitive-based models^12^, only in recent years muscle synergies analysis has been used to describe adaptive behaviors, including visuomotor adaptations during isometric reaching movements^35-37^. In this latter scenario, muscle synergies have been shown to be robust with respect to changes in the motor plan^36^, and that adaptation to specific perturbations depends on how the perturbations map on the synergistic structure^35,37^.

If adaptation to visuomotor rotations depends from the rotation of synergies of specific shape and workspace, it is left to understand if such synergies rotate solidly and invariantly in response to a consistent perturbation or if this rotation depends on the spatial and temporal characteristics of the training. In addition, synergies rotation could also better explain adaptation-related phenomena such as generalization. Generalization to visuomotor rotations has, in fact, been observed to depend on both the distance between the pre-adapted targets and the newly experienced ones^38^ and the amount of angular rotation of the perturbation that needs to be generalized^39^. In a synergistic model, both phenomena could depend on how specific synergies rotate across the workspace.

Here we studied (1) if synergies always rotate in the same way when adapting to a visuomotor rotation and (2) if synergies rotation can explain generalizing behaviors. We hypothesized that the presence and extent of synergies rotation inherently depends on the way in which the adaptation is presented (specifically, the order of the targets that are adapted) and that generalization and non-generalization behaviors strictly depend on the rotation of specific synergies during the first set of adaptation.

**Figure 1.**
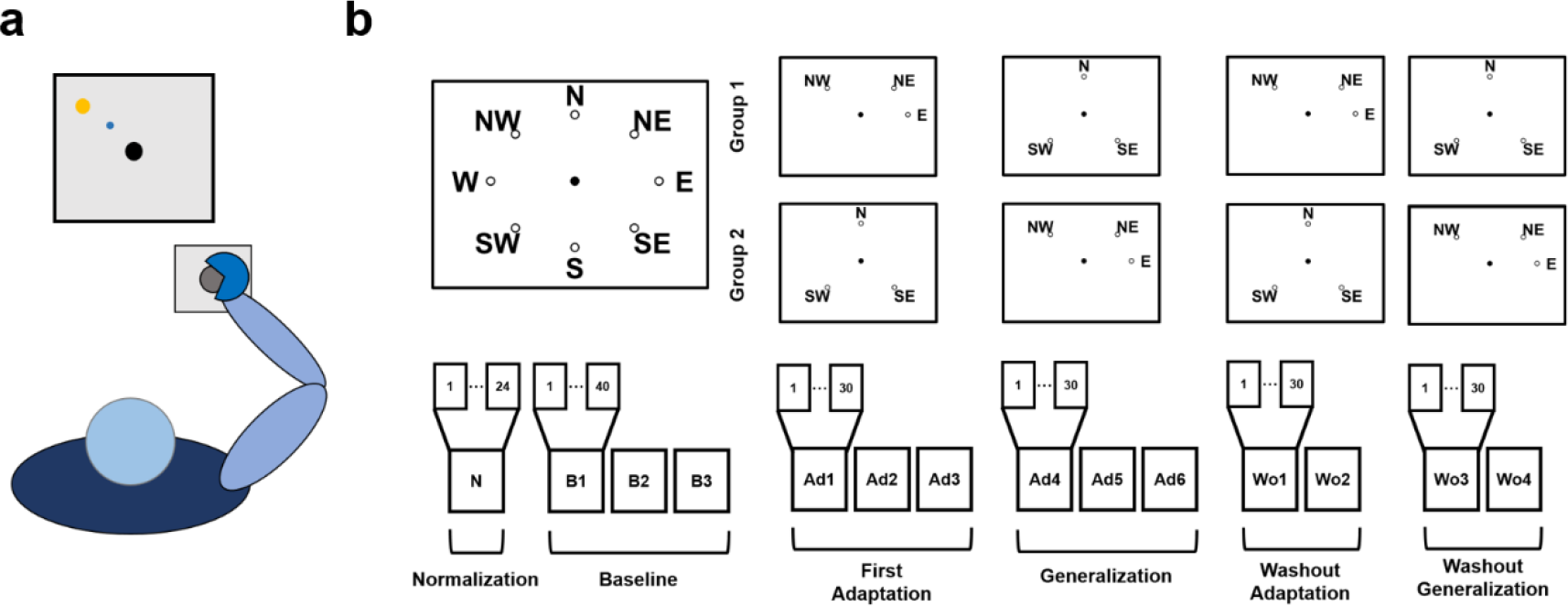
The experimental setup and protocol. Panel (**a**) presents a graphical representation of the task. Subjects kept their position consistent across all trials. The forearm was strapped to a support surface (not-shown in the picture) and the hand was strapped to the manipulandum to avoid the use of the hand muscles during the task. Subjects were presented a virtual scene on a screen in front of them. During the whole experiment, subjects performed a total of 14 exercise blocks (**b**). The number of movements experienced in each block varied between 24 and 40. During the normalization (N) and baseline (B1-B3) blocks both groups of subjects experienced all 8 targets (3 repetitions of each target during normalization, 5 during baseline). During the first adaptation (Ad1-Ad3) and generalization (Ad4-Ad6) blocks subjects experienced a limited target-set of 3 targets (NW, NE and E for target-set 1; N, SE and SW for target-set 2; each target was experienced 10 times in each block). Group1/Group2 experienced target-set 1/target-set 2 during first adaptation and target-set 2/target-set 1 during generalization. A CW visuomotor rotation of 45° was applied during Ad1-Ad6. During the washout blocks (Wo1-Wo4) subjects experienced the same target-sets as in Ad1-Ad6, without the visual rotation.

## Results

### Experimental Protocol

We designed a single experiment that could be used to test our initial hypotheses (Figure 1). In the experiment, subjects trained to counteract a 45° clockwise (CW) rotation during isometric reaching movements. Subjects first experienced a series of baseline blocks (B1-B3) where they practiced reaching to eight different targets positioned in a compass-like configuration and at a distance equivalent to 15 N. After that, subjects trained to adapt six of the eight available targets to the CW rotation, divided in two target-sets of three targets each. The subjects experienced one of the two target-sets during three blocks of adaptation (Ad1/Ad3) and the other one during three blocks of generalization (Ad4//Ad6). The subjects were divided in two groups: Group1 adapted target-set 1 (TS1 that included targets NW, NE and E) during the adaptation blocks and target-set 2 (TS2 that included targets N, SE and SW) during the generalization blocks, while Group2 adapted TS2 during the adaptation blocks and TS1 during the generalization blocks. After the generalization blocks subjects performed two sets of washout blocks (two blocks per set) where they first washed out the target-set adapted during the adaptation phase (Wo1/Wo2) and then that adapted during the generalization phase (Wo3/Wo4). TS1 and TS2 were designed so that one target-set would span only a limited part of the workspace (TS1, spanning a total of 135°) while the other one would present targets scattered through all the workspace (TS2). This choice was made under the hypothesis that training on a smaller sub-space would lead to the rotation of a smaller sub-set of synergies with respect to a wider sub-space, thus allowing to test the hypothesis of differential rotation. In addition, this choice of targets allows for different configurations between pre-adapted and generalized targets. Specifically the two target-sets include: i) targets with no adjacent pre-adapted targets (SW, TS2); ii) targets with one pre-adapted target (NW, NE and E in TS1 and SE in TS2); iii) targets with both adjacent targets pre-adapted (N, TS2).

### Adaptation and generalization in the biomechanics

The analysis of the force trajectories recorded during the experiments showed that: a) subjects were able to fully adapt to both target-sets regardless of whether they were presented during adaptation or generalization; b) the marked patterns of generalization that can be observed across the different targets are generally dependent on the distance between the generalized targets and the previously adapted ones. These results are presented in Fig. 2.

**Figure 2.**
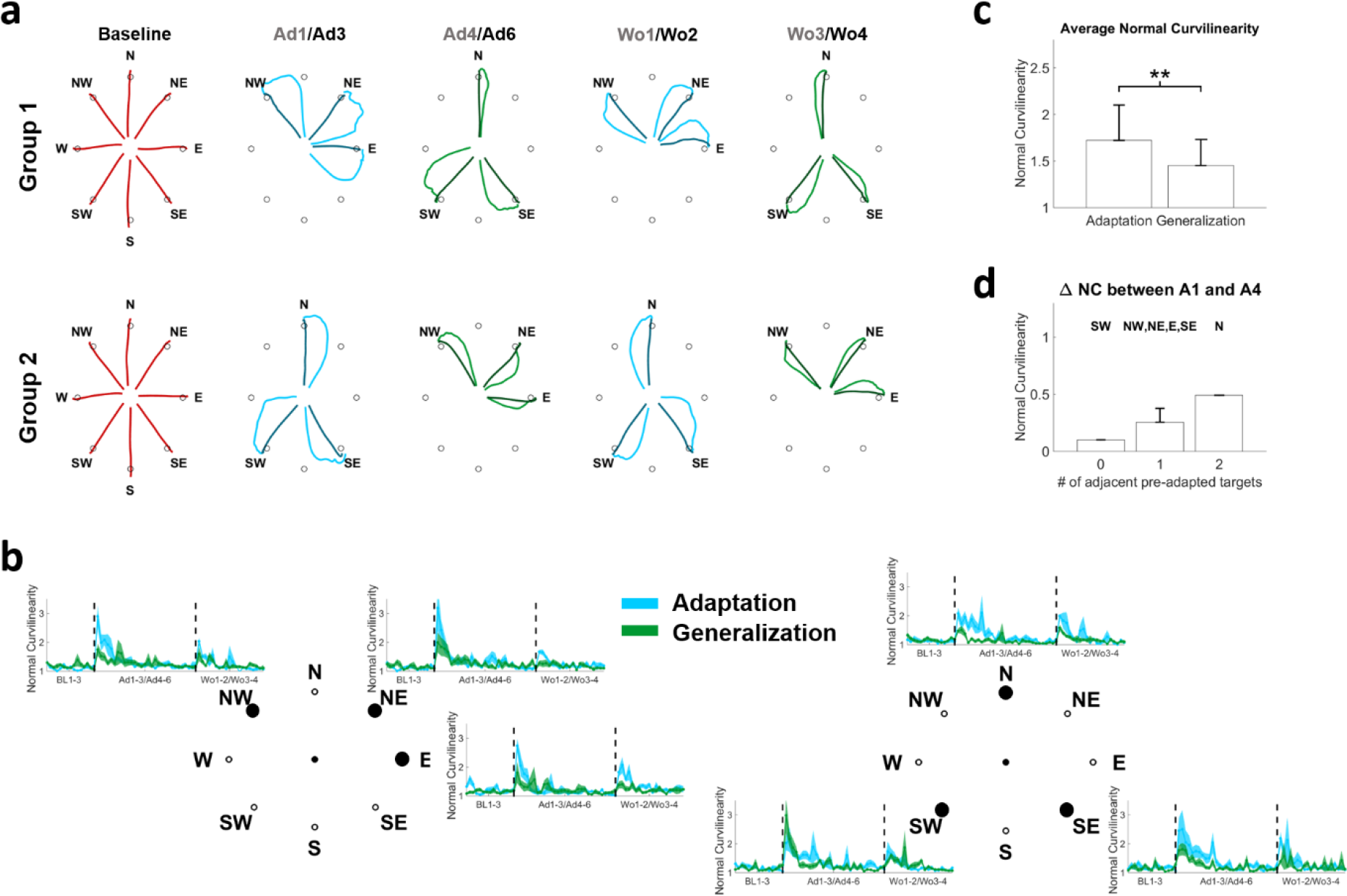
Adaptation and generalization in the biomechanics. Panel **a** presents the average trajectories observed at Baseline (red), Ad1/Ad3 (blue), Ad4/Ad6 (green), Wo1/Wo2 (blue) and Wo3/Wo4 (green). For each plot the lighter color presents the average of the first 2 movements in the first block listed on top (e.g. Ad1) and the darker color presents the average of the last 2 movement in the second block listed on top (e.g. Ad3). For baseline, the average of the last 5 movements of B3 is presented. For each target we observed an increase in movement curvilinearity at the onset of the perturbation (**b**) followed by an adaptation and a second increase when the perturbation is removed. Similar behaviors were observed during adaptation (light blue) and generalization (light green) although generalization was characterized by smaller initial deviations at the onset and offset of the perturbation for all targets with the exclusion of target SW. The average normal curvilinearity was significantly higher at the beginning of adaptation with respect of generalization across all targets (**c**). ** denotes p<0.01, using Mann-Whitney U test. Panel **d** shows the average values of generalization (calculated as the average difference in NC between Ad1 and Ad4 across the subjects) for the targets as a function of the number of pre-adapted adjacent targets.

As expected, both groups presented an effect of the visual perturbation at the beginning of the adaptation period that was characterized by a CW-rotated initial trajectory followed by a compensatory movement (Figure 2a). Subjects were able to quickly and fully adapt to this perturbation and were able to perform the movements with straight trajectories by the end of the adaption period. At the beginning of the first generalization block, both groups presented smaller initial deviations that were again fully compensated. After the visual rotation was removed, subjects presented marked after-effects during the washout periods of the adaptation target-set and, to a lesser extent, of the generalization one. We analyzed the patterns of adaptation and generalization in terms of the Normal Curvilinearity (NC) metric that is calculated as the ratio between the length of the actual and ideal trajectories. This metric was selected because, with respect to other metrics such as the initial angular error^36^, it keeps into account both the initial angular error and potential compensatory movements. As preliminary observed, subjects presented values of NC close to 1 (equal to the ideal straight trajectory) during baseline (Figure 2b). Once the visual perturbation was introduced (either during adaptation or generalization) the value of NC suddenly increased and was then reduced again close to 1 by the end of the last adaptation/generalization block (Ad3/Ad6). Once the perturbation was removed, subjects once again exhibited an increase in NC (indicating the presence of the after-effect) that was quickly washed out. For all targets, with the exclusion of SW, the NC values recorded during the first block (10 repetitions) of generalization were generally lower than the respective values during adaptation. For all targets both groups showed the same level of final adaptation/generalization in the last perturbed block (Ad3/Ad6). Similarly, after-effects were shown to be lower for generalization than for adaptation in the first washout block for all targets with the exclusion of SW. These results suggest a marked presence of generalization in our tests, characterized by a pre-learning effect of previous adaptation to targets in a shared workspace on targets that have not been trained yet. To confirm upon this point we compared, across all subjects, the average NC calculated for each target during the first adaptation and first generalization blocks, for all targets pooled together. We observed (Figure 2c) a statistically significant effect of pre-adaptation where subjects presented, over the same targets, a bigger biomechanical error during the first block of adaptation with respect to the first block of generalization (NC = 1.72±0.38 for adaptation, 1.45±0.28 for generalization, p<0.01 using Mann-Whitney U test). Finally, in order to check for a dependency of generalization from the distance to pre-adapted targets, we calculated the extent of generalization for each target as the difference between the average (across repetitions and subjects) NC during the first block of adaptation and the first block of generalization and grouped the targets depending on how many adjacent pre-adapted targets they had during generalization. The smaller amount of generalization was observed for target SW (Figure 2d) that was the only target that did not have adjacent pre-adapted targets. Targets that presented one adjacent pre-adapted target showed limited generalization. The only target that had both adjacent targets pre-adapted before Ad4 (target N) showed the highest level of generalization.

### Adaptation is achieved through the rotation of fixed motor modules

Gentner and colleagues^36^ have shown that adaptation to visuomotor rotations is fully explained by a rotation (or tuning) of the activation patterns of a fixed set of muscle synergies. Here we demonstrate that the motor module themselves do not change during this kind of adaptation, thus the synergies recorded at baseline can be used to reconstruct the muscular activity recorded during the whole experiment. Similarly to previous works^36,40^, we found that four muscle synergies, extracted using the non-negative matrix factorization algorithm (NMF)^41^ could satisfactorily reconstruct the EMG activity of all subjects during baseline (average Variance Accounted For (VAF) across subjects = 91±2%).

**Figure 3.**
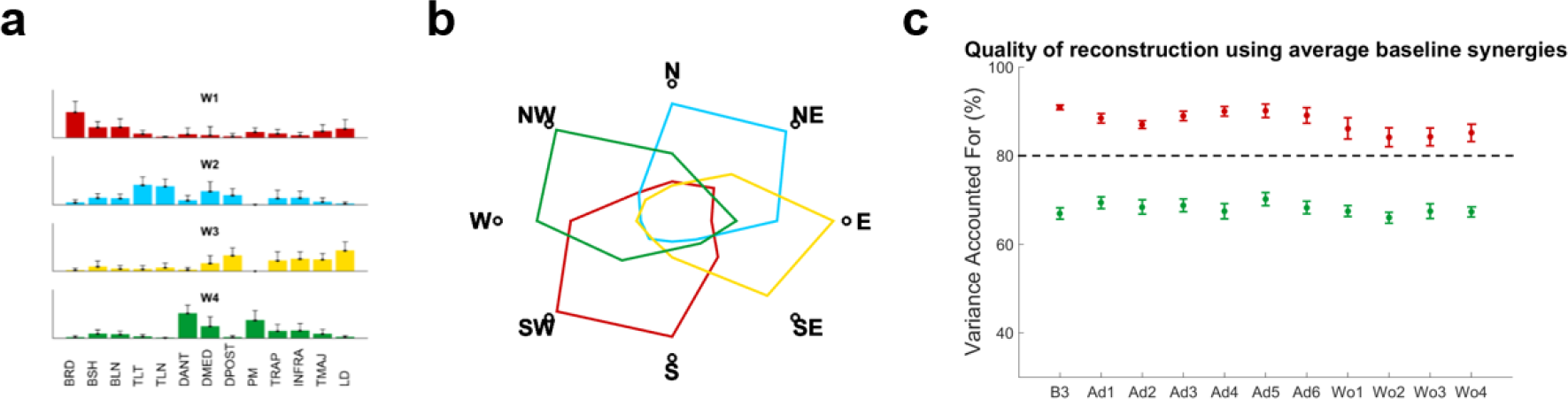
Muscle Synergies extracted at baseline. Four muscles synergies could well represent the whole dataset (**a**). The RMS values of the activation patterns of each synergy at each target were used to represent the spatial boundaries of each module in the workspace (**b**). The modules (**a**) extracted during B3 were used to estimate the activation patterns able to describe the movements in all the blocks. We found that the four original synergies were able to represent the data with acceptable median VAF (>80%) in each block (**c**). The VAF calculated for each block (red errorbars, indicating median and standard error) were significantly higher than the ones obtained using random modules (green errorbars). For each block statistical significance was estimated using Mann-Whitney U test and Bonferroni’s correction.

These four synergies (*W1-4*) were shown to have distinct functional characteristics and span specific sections of the experimental workspace (Figure 3a-b). *W1* is constituted by the elbow flexors and acts mainly in the lower-left quadrant of the workspace. *W2* is constituted by the elbow and shoulder extensors and acts mainly in the upper-right quadrant. *W3* is constituted by the muscles commanding shoulder extension and act mainly in the lower-right quadrant. *W4* is constituted by muscles involved in shoulder flexion and act mainly in the upper-left quadrant. In our synergistic model of adaptation we described the adaptive behavior as the modification of the activation patterns (APs) of a fixed set of reference muscle synergies. For each subject, we fixed the reference synergies to those extracted in the last block of baseline (B3), chosen as the block where subjects were fully habituated to both the task and the movement environment. We then reconstructed the APs relative to the fixed synergies for each block using a modified version of the NMF algorithm^40^. The product between the reference synergies and the block-specific APs yielded average values of VAF>80% across all subjects and were shown to be significantly higher (p<0.0045 for each block, using Mann-Whitney U test and Bonferroni’s correction) than VAF values obtained using random reference synergies (Figure 3c).

### Differential paths to adaptation

We then analyzed how the synergies APs converge at the end of adaptation/generalization. Figure 4 shows the average shape of the APs of each target during the last 5 movements of Ad3/Ad6. Since the visual perturbation consists of a 45° CW rotation, in order to adapt and perform a straight trajectory subjects need to reach for each target using the motor plan they would use for the adjacent 45° CCW-rotated one. This corresponds to a 45° CW rotation of the synergy APs. As a general observation, the adapted shapes of the APs roughly resemble those observed at baseline for the 45° CCW-rotated target. The average across targets similarity between the adapted APs and the CCW-rotated ones (calculated as the correlation coefficient between the average of the last five movements at Ad3/Ad6 and the last five movements of the adjacent CCW target at B3) is equal to 0.59±0.15 for adaptation and 0.61±0.06 for generalization, while the similarity between the adapted APs and the baseline ones for the same targets is equal to 0.38±0.17 for adaptation and 0.45±0.13 for generalization. Thus, after adaptation, the APs resemble more those of 45° CCW-rotated targets than the original ones. In Fig. 4 we also notice that the adapted APs differ, for some targets, between the group that trained that target during adaptation and the group that did it during generalization. This behavior can be observed clearly in the APs of *W1* for target SW, *W3* for targets NE, E and SE and *W4* for targets E and SE. This qualitative observation suggests that the synergies APs rotate differently between the two groups.

**Figure 4.**
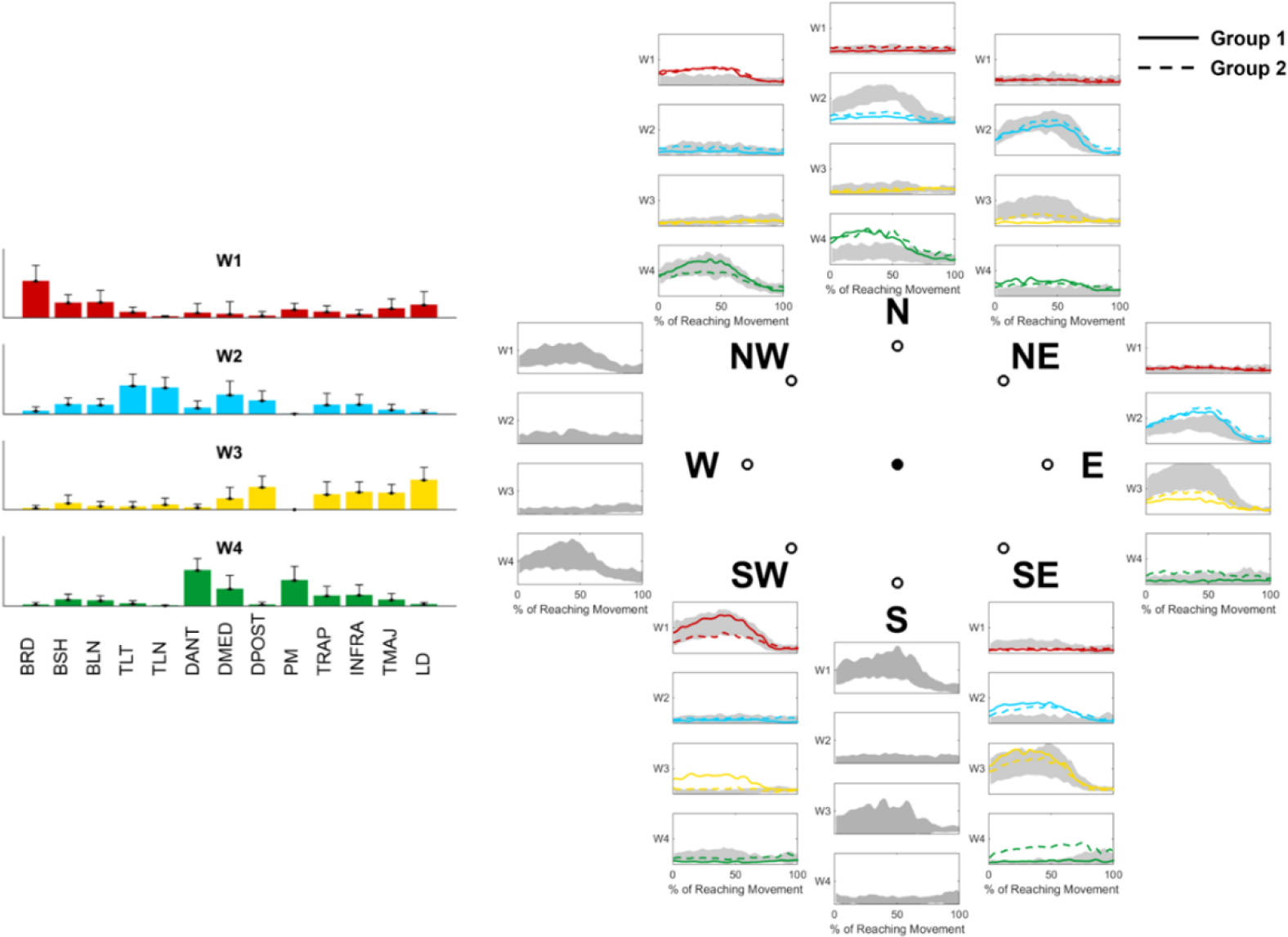
Synergies APs at baseline and final adaptation for each target. Each plot presents the activation patterns (color-coded as their relative synergies on the left of the plot) of each synergy AP for each target. Grey shades represent the average and standard deviation across all subjects at B3, solid lines present the average of the last 5 repetitions of Ad3/Ad6 for Group1, while dashed lines represent the average of the last 5 repetitions of Ad3/Ad6 for Group2.

To confirm upon this point, we calculated and compared the preferred directions of the synergies APs at baseline and after targets were fully adapted (either during adaptation or generalization) for both groups (Figure 5). The preferred direction of each AP was calculated from the coefficients of the cosine fitting of the average (across subjects) AP intensity values of all targets and the corresponding target positions^28,36^. For each group, targets trained during adaptation and generalization were pooled together while calculating the cosine fitting relative to full adaptation. Thus this analysis does not take into account possible modifications in the APs of the targets trained in Ad1/Ad3 that happen during the Ad4/Ad6 blocks. This analysis can then be considered as a snapshot of the tuned APs relative to each target at their first instance of full adaptation. The comparison between the polar representations of the baseline and adapted APs activation intensities (Figure 5, first row) confirm the hypothesis that adaptation is achieved by tuning the APs by rotating them CW. However, evident differences can be observed between the two groups, especially for *W1* and *W3*. The analysis of the preferred angles (Figure 5, second row) confirmed this observation. Group1 presented rotations equal to 24°, 56°, 65° and 36° for *W1-4*, while Group2 presented rotations of 43°, 43°, 27° and 50°. Interestingly, Group2, that first trained on the target-set spanning the whole workspace (TS2), presented values of final rotations close to the expected 45°, while Group1 presented over-rotation for two of the synergies (*W2* and *W3*) that cover the workspace of TS1. It appears clear then that the two groups, although presenting the same biomechanical errors at the end of the adaptation/generalization phases of all targets, achieve full adaptation in different ways.

**Figure 5.**
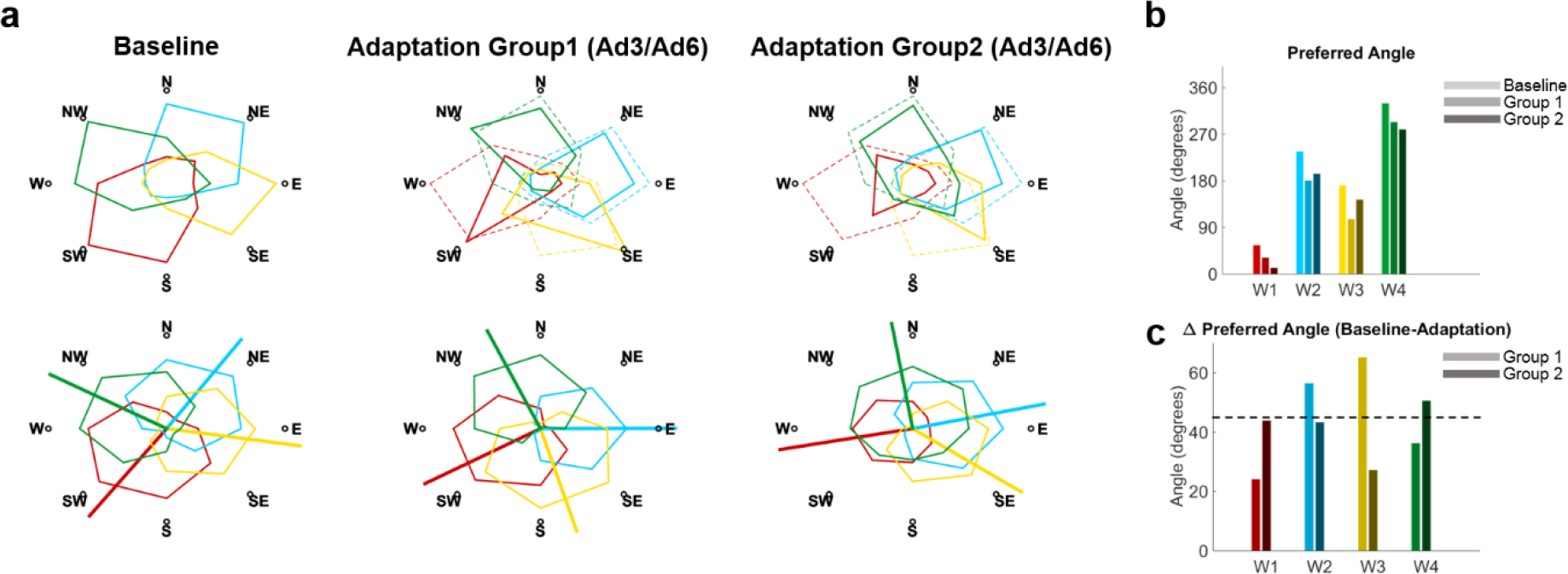
Rotation of the synergies activation patterns and preferred angles. The first row of panel **a** shows the average RMS of the synergies activation patterns (color coded as in Fig. 4) across all targets during Baseline and Full Adaptation for both groups. The Full Adaptation plots were obtained by pooling together the RMSs of the targets at the end of Ad3 (end of first adaptation) and Ad6 (end of generalization). Dashed lines represent the Baseline synergies rotated 45° CW. Both groups present rotations consistent with a 45° CW rotation of the synergies at baseline. The second row presents the same data interpolated using a cosine fit. Outward-pointing lines represent the preferred angles (estimated from the fitting parameters) of each synergy. We observed changes in the preferred angles between Baseline and Full adaptation across the two groups (**b** and **c**). Most preferred angles across the four synergies presented rotations close to the expected 45° (**c**). The biggest differences in preferred angles after adaptation between the two groups were observed for W1 and W3.

We then analyzed the path to full adaptation for both groups from baseline to washout (Figure 6). Group1 was able to quickly tune the APs of *W2* and *W4* at the beginning of Ad1 and by the end of Ad3 presented an over-rotation of *W2* (suggested by the decreased activation at target E) and small rotations for both *W1* and *W3* (characterized by an increase activation of *W1* at NW and by the almost disappearance of *W3* from the adapted workspace of TS1). Thus at the end of adaptation, Group1 had tuned, to different extents, all four synergies. At the beginning of generalization (Ad4) the four synergies APs appear to be already tuned as highlighted by their similarity with the baseline-rotated ones. By the end of Ad6, Group1 appeared to have further tuned *W3* CW (as suggested by the increase in activity in SW) while the increased activity of *W1* at target SW suggests a small CCW counter-rotation for that synergy. At the beginning of Wo1, Group1 quickly de-tuned all APs close to their initial state, which was mostly reached by the beginning of Wo3 although with a residual rotation effect on all synergies. As a summary, Group1 was able to tune all four synergies by rotating them CW during the exposure to TS1 (Ad1-Ad3), with an emphasis on the synergies mostly spanning the workspace (W2, W3 and W4). During generalization, only small adjustments to the adapted state were sought and regarded exclusively target SW that is the only target that, in our results, did not present a clear generalization behavior.

Group2 reacted to the onset of the perturbation at Ad1 by quickly rotating *W1* and *W4* and, to a lesser extend *W2* that was fully tuned only at the end of Ad3. *W3* appeared to be only slightly rotated during the adaptation period (Ad1-Ad3), most likely because the activity of this synergy is very similar between SE and E (that is SE’s adjacent CCW target, thus its goal-target for adaptation). This observation is confirmed by the very limited tuning exhibited by *W3* at the beginning of Ad4 while *W4*, on the other hand, appeared to be slightly over-rotated. At the end of Ad6 all 4 synergies presented a small additional CW rotation with respect to Ad4. At the beginning of the first washout phase *W3* appears to be quickly de-tuned, while *W1*, *W2* and *W4* still present residual tuning. Finally, at the beginning of Wo3 only the *W4* synergy appeared to be still rotated. As a summary, Group2 only tuned three of the four synergies during the exposure to TS2, while subsequent adaptation to TS1 was achieved by rotating the previously non-tuned synergy (*W3*) and only slightly adjusting the rotation of the other ones. Taken all together, the results presented in this section suggest a dependency of final adaptation on the way in which the synergies are tuned during the initial adaptation period.

**Figure 6.**
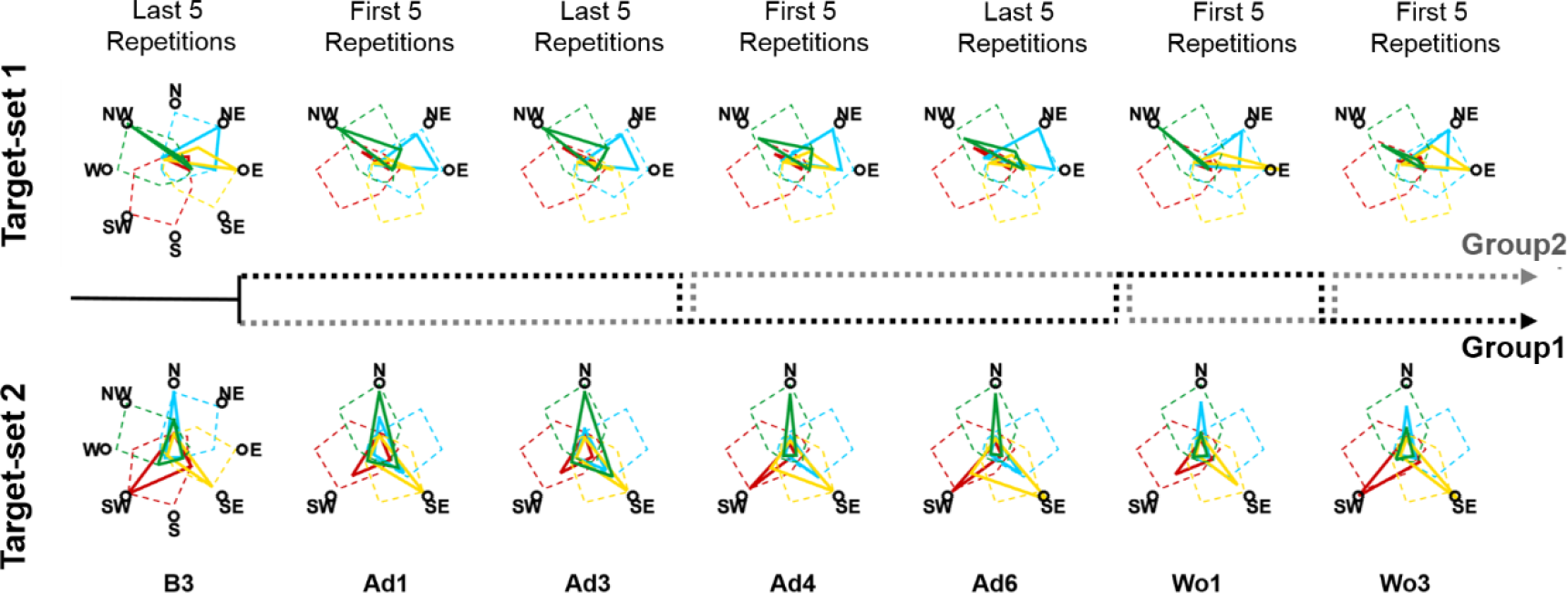
Differential paths to adaptation. The top row presents the changes in AP intensities for TS1 across the different blocks, while the bottom row presents the same results for TS2. For blocks B3, Ad3 and Ad6 the solid lines (color coded as the synergies in Fig. 4) represent the average intensities of the last 5 repetitions for each target, averaged across subjects, while for blocks Ad1, Ad4, Wo1 and Wo3 the solid lines represent the average intensities of the first 5 repetitions for each target, averaged across subjects. For the B3 plots the dashed colored lines represent the workspace spanned by the synergies at baseline as reconstructed using all the targets, while for all the other blocks the dashed colored lines represent the workspace spanned by the synergies at baseline rotated 45° CW. The dashed black and grey lines between the two rows represent the order of repetition for each group. Group1 (black dashed line) performed TS1 during Ad1-Ad3, TS2 during Ad4-Ad6, TS1 during Wo1 and TS2 during Wo3. Group2 (grey dashed line) performed TS2 during Ad1-Ad3, TS1 during Ad4-Ad6, TS2 during Wo1 and TS1 during Wo2.

### Generalization in the synergistic representation of movement

We hypothesized that, if the movement space and thus the adaptation behavior are bound to the workspace spanned by the existing muscle synergies^35-37^, generalization behaviors will also depend on how the synergies are pre-trained. In our experiment we observed patterns of generalization consistent with this hypothesis. The target presenting the higher degree of generalization is target N, which is the only target that, during generalization, had both adjacent targets already pre-adapted. Group1, which is the group that experienced target N during generalization (Figure 7a), presents shapes of APs during the first 5 movements of generalization that are remarkably similar to those observed in the baseline CCW-rotated target. We observed a significant (p<0.01, rho=-0.34) linear correlation between the values of NC and the similarity between the APs at Ad1/Ad4 and the baseline CCW-rotated target (Figure 7b). We also observed that similarity during generalization is significantly higher than during adaptation (Figure 7c, p=0.03). These results taken together further confirm that adaptation and generalization to a CW visuomotor rotation depend on a CW tuning of the synergies.

To better understand the reasons behind the different levels of generalization that we observed across the different targets, we analyzed the patterns of generalization with respect to the way in which the synergies are tuned for each target in both groups (Figure 7d). Target SW, that is the only target that does not present generalization, is adapted by tuning *W1* and *W3* that are the two synergies presenting the bigger differences in rotation after adaptation between the two groups (Figure 5c). Target N, on the other hand, is the only target whose adaptation depends on the tuning of synergies *W2* and *W4*, which are the two synergies showing the closest preferred angles between the two groups after adaptation. Plotting the average APs similarity with the baseline CCW-rotated targets over the difference in preferred angles calculated between the two groups, it is possible to appreciate a trend where targets associated with synergies showing higher differences in the preferred angle after tuning present lower overall similarity (Figure 7e). Nevertheless, the differences across the four groups presented in Fig. 7e were not significant (p = 0.09 based on Kruskal-Wallis test). These results, although qualitative, suggest that generalization is synergy-dependent and correlates with how similarly the APs tune, during the first adaptation period, towards the optimally tuned preferred angles.

**Figure 7.**
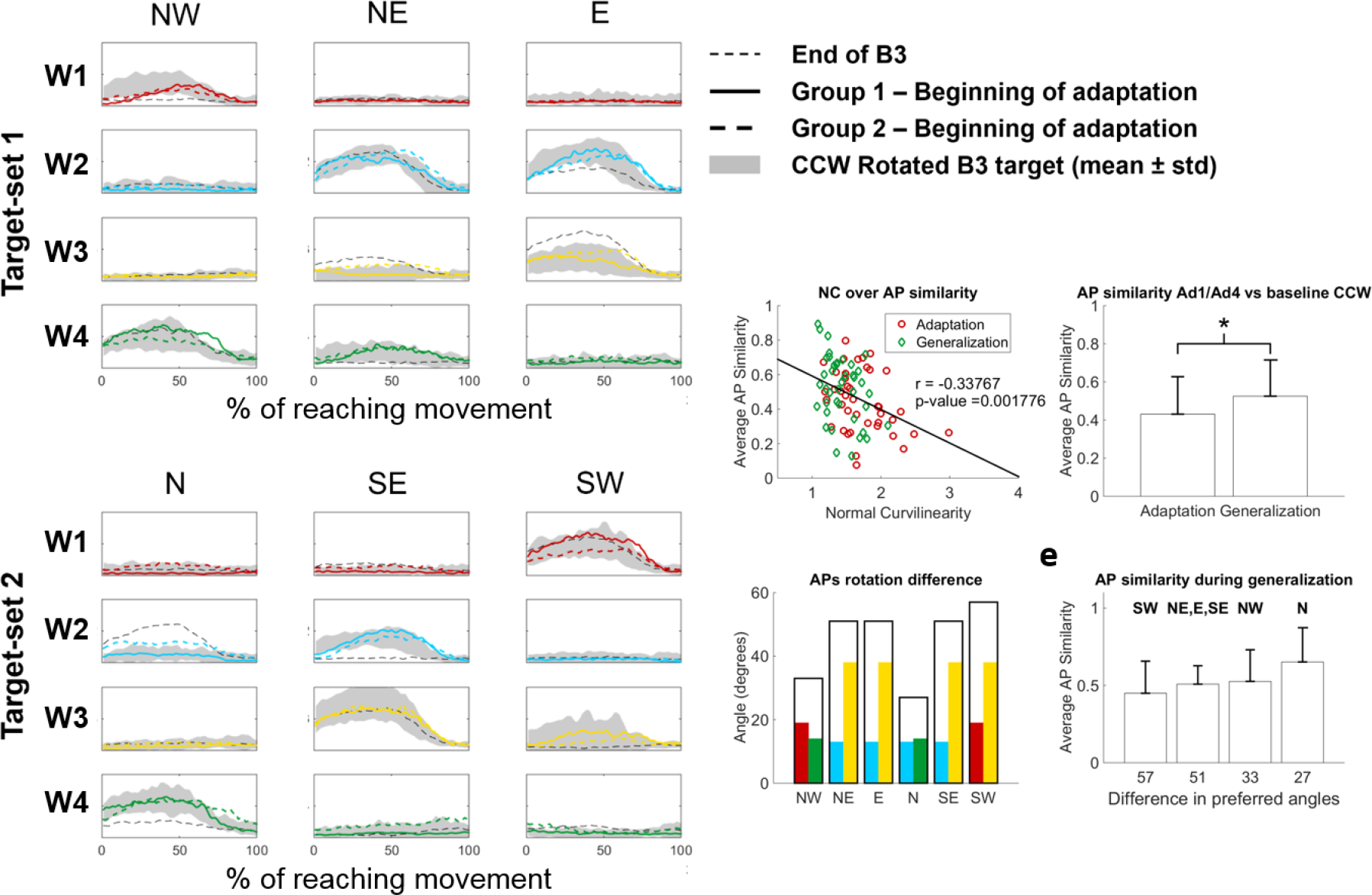
Generalization in the synergies APs. Panel (**a**) shows the APs of the four synergies (color coded as in Fig. 4) at the beginning of Ad1/Ad4. Solid lines are relative to Group1, while dashed lines are relative to Group2. Shaded areas in each plot represent the activity of each AP in the 45° CCW target at B3 that theoretically represents the goal AP activity for adaptation. The black solid line represents the activity of the APs for each target at the end of B3. For the group/target combination that presents the higher level of generalization (Group1, target N, solid line) the APs at the beginning of generalization are perfectly consistent with the baseline CCW-rotated ones. We observed a significant correlation between the similarity of the APs at the beginning of Ad1/Ad4 with the baseline CCW rotated ones and the values of NC (**b**) indicating that adaptation and generalization in the biomechanical metric correlates with a 45° rotation of the synergies AP. This result is also confirmed by the fact that the similarity of the APs with the baseline CCW rotated ones is significantly higher for Ad4 with respect to Ad1 (**c**) (p=0.03, Mann-Whitney U test). We observed a qualitative correlation between the difference in preferred angles at adaptation for the two groups and the level of generalization (**d-e**). Specifically, the target presenting the higher level of generalization (both in terms of NC and synergies similarity with the baseline rotated ones) is also the target that adapts by tuning the synergies that are more similar between the two groups.

## Discussion

Here we have shown that adaptation to visuomotor rotations can be achieved through different rotations of the same set of muscle synergies. The two different final tunings depend primarily on the initial rotation achieved when adapting to a first sub-set of targets, while subsequent exposures to untrained targets appear to lead only to small adjustments that happen mostly in un-trained or less-trained parts of the workspace. We also show that there is an influence of synergistic structure on the presence and extent of generalization after pre-adaptation to targets in the same workspace.

Rotational tunings in muscular and M1 activities have been observed to happen during motor adaptations to visual rotations and force-fields^42-44^. Those results have been so far extended also to muscle synergies^35,36^, by demonstrating that simple visuomotor rotations, compatible with the existing synergies workspace, are achieved rapidly through tuning of synergies APs, while more complex adaptations non-describable using the already present synergistic structures may require a slower rewiring of the synergies themselves^12,35^. The fast rates of adaptation observed in our results suggest that only the former component of adaptation is at play in our experiment.

We observed, as initially hypothesized, differential paths of APs tuning for the two groups, characterized by under-rotations (*W1* for Group1, *W3* for Group2), over-rotations (*W2* and *W3*, Group1) and theoretically-optimal rotations (*W4* for Group1, *W1*, *W2* and *W4* for Group2). Interestingly, most examples of optimally tuned synergies have been observed in Group2, where three of the four synergies appear rotated close to the expected 45°. It is possible that this result is correlated with the fact that Group2 first trains on the target-set (TS2) that spans all the workspace. On the other side, Group1, which was initially trained on a smaller sub-space, presents non-optimal tuning in three of the four synergies. These results, taken together, suggest that optimal synergies tuning is obtained through optimal sampling of the movement space.

A remarkable result herein presented is that it is not enough for a synergy to span the reaching workspace of a given target in order for it to be tuned in response to a visuomotor rotation. This is exemplified by the behavior that we observed for synergy *W3* in Group2. Synergy *W3* spans the workspace of targets NE, E and SE, and, when tuned in response to a CW 45° rotation, that of targets E, SE and S (Figure 5a). TS2 only engages *W3* for the adaptation of target SE. Nevertheless, since the activity of *W3* is the same between targets SE and E (that is, in this case, the tuned version of target SE), Group2 (that experience TS2 during the first adaptation) does not tune *W3* during the first adaptation period, as confirmed by its non-rotation at Ad4, and instead adapts target SE by tuning *W4* (Figures 4 and 6). Group1, on the other hand, tunes *W3* while adapting for target E, whose goal-target NE presents a completely different AP shape for *W3*, and uses the tuned synergy also when adapting to SE during the generalization trials. The only other example of an AP that does not necessarily require tuning for adaptation to a target is, in our experiments, *W2*, when tuned to adapt for target NE. *W2* presents in fact the same activation profile for both NE and its goal-target N. Nevertheless, Group1 still tunes *W2* during the adaptation trial likely because this synergy is also engaged for the adaptation of target E, which requires a modification in its AP. The results on *W3* and *W2* seem to indicate that synergies are tuned only if engaged at the boundaries of their workspace.

The patterns of differential adaptation that we observed suggest that the synergies APs are not the parameters that are directly adapted by the CNS, but rather the tools that are differentially tuned to adapt, while high-level adaptation is likely obtained by updating the sensorimotor transformations mapping the visual targets to the force directions^36,45^. Then, as different muscular activation patterns can produce the same force^46^, similarly can different combinations of synergies APs, without necessarily invalidating the synergies hypothesis. From our results it appears that muscle synergies constraint the adaptable mapping between the desired motion and the motor commands^2^. Given the sensory prediction error calculated in the forward representation of movements^47^, motor commands are then updated in the synergy-space in a parsimonious way, thus leaving un-altered synergistic structure (unless necessary)^35^ and APs compatible with the desired force output.

We initially hypothesized that, if adaptation patterns are constrained by the synergistic structural components of the workspace (as our experiment appears to confirm), this likely reflects also on how training on an initial set of targets generalizes on a set of newly experienced ones in the same workspace. In accordance with our hypothesis, the results of our experiment show that the patterns of generalization after pre-adaptation depend on pre-tuning of specific synergies. The target presenting the higher level of generalization is target N, which requires the tuning of synergies *W2* and *W4*. As observed in Fig. 6 both groups are able to engage these two synergies at their boundaries (Group1 while training NW, NE and E; Group2 while training N and SE) in two sub-spaces of identical extension (135°) and rotated of only 45° among them. As a result, the final tuning of those two synergies is similar between the two groups and close to the theoretically ideal rotation (45°). The biggest differences in AP tuning between the two groups are observed in *W3* and, to a smaller extent, *W1*. It is then interesting to notice that all the targets showing limited generalization recruits at least one of these two synergies for its adaptation (Figure 7d), while target SW, that does not show generalization, recruits both.

Our results are in agreement with previous studies showing that generalization to visuomotor rotations depends on the angular distance between the pre-adapted targets and the new ones^16,38^. In fact, in the synergy-tuning model of adaptation, pre-training on sub-spaces close to the one that needs to be generalized is likely to lead to the adaptation of one or more of the synergies that are involved in the adaptation of the new movement. This conceptual model could also explain the results showed by Abeele and colleagues^39^, who found that visual rotations up to 120° present a beneficial effect of pre-adaptation to a 45° smaller rotation, but rotations bigger than 120° did not present a beneficial effect of pre-training. These behaviors may depend, at the synergistic level, on the discrepancy between the original synergies, the pre-tuned ones and the ones that need to be ultimately tuned. Our results suggest that if the synergy sets involved in all three movements (original, pre-adapted and generalized) fully overlap, generalization is maximized, while as different synergies are recruited (as it is likely for big deviations, as in our 4-synergy space most movement are driven by a couple of APs that usually span about 160°) generalization is reduced or lost.

In conclusion, our results present evidence that the modularity in motor commands as explained by a muscle synergies model influences the generation and generalization of motor adaptations. In light of the rich literature describing visuomotor adaptations, our results present the rationale for taking into account the functional constraints posed by modular organization of movement in the interpretation of the results of adaptation experiments. In this vision, muscle synergies may represent the functional primitives (or a part of) that are updated in the inverse representations of the motor system during motor adaptations. The link between muscle synergies, adaptations and, possibly, the mechanisms of motor learning, could have sensible practical validity in the field of neurorehabilitation, where synergistic representation of the impaired and target movements, and of the “modular” path that can be used to translate the former into the latter, could be used to design ad-hoc therapies for patients.

## Methods

### Experimental setup and Protocol

Fourteen healthy individuals (7 females, age 25.6 ± 1.2) participated in this study. Each individual participated in a single experimental session consisting of a series of isometric reaching tasks performed with their right arm. All the experimental procedures describe in the following have been approved by the Ethical Committee of University College Dublin and has been conducted according to the WMA’s declaration of Helsinki. All subjects gave written informed consent before participating in this study. During the exercises the subjects sat in a chair with their back straight. Their right forearm was put on a support plan. The hand was strapped to a fixed manipulandum (consisting of a metal cylinder of 4 cm of diameter) attached to a tri-axial load cell (3A120, Interface, UK), while the wrist and forearm were wrapped to the support plan and immobilized using self-adhesive tape. Data from the load cell were sampled at 50 Hz. During all exercises, subjects kept their elbow flexed at 90° and their shoulder horizontally abducted at 45° (Figure 1a), so that the manipulandum would be exactly in front of the center of rotation of their shoulder. The elevation of the chair was controlled so to keep the shoulder abducted at 100°. Subjects sat in front of a screen displaying a virtual scene. The virtual scene consisted of a grey cursor, commanded in real time by the *x* and *y* components of the force exerted on the load cell through the manipulandum, a filled circle indicating the center of the exercise space and, depending on the phase of the exercise, a target, represented by a hollow circle. Both the center and target circles had a radius of 1.3 cm. Across all the blocks of the experiment subjects experienced a total of 8 different targets, positioned in a compass-like configuration (Figure 1b) at a distance of 9.5 centimeters from the center of the screen. The force-to-pixel ratio of the virtual scene was set so that each target was positioned at a distance of 15 N with respect to the center (representing 0 N applied to the load cell) of the scene. The virtual scene and the exercise protocol were controlled using a custom software. The experiment was divided in fourteen different blocks during which subjects were asked to reach for different targets shown on the virtual scene. Between different blocks or different repetitions of the same block subjects were allowed to rest for 1 minute.

At the beginning of the experiment subjects performed a Normalization (NM) block. In this block subjects were asked to reach for each one of the eight targets three times and hold the cursor on the target for 5 seconds. In this and all subsequent blocks the targets were presented in a pseudo-random order with the constraint that the same target could not be shown consecutively more than two times. The EMG signals recorded during this block were used to normalize the amplitude of the EMGs for the estimation of muscle synergies. After the NM block, subjects performed three Baseline (B1-B3) blocks. In these blocks subjects were asked to reach for each of the eight targets five times in a random order. In these and all subsequent blocks subjects were asked to complete each reaching movement in less than 1.5 seconds, calculated between the instant in which the cursor exited the central mark circle up to the instant it reached the target hollow circle. Subjects were shown positive feedback for movements performed in less than 1.5 seconds, consisting in the target circle turning green, and negative feedback for movements performed in more than 1.5 seconds, consisting in the target circle turning red. Targets that were performed too slowly were discarded and repeated at the end of the block. After the B1-B3 blocks, subjects performed three Adaptation (Ad1-Ad3) and three Generalization (Ad4-Ad6) blocks. In all these blocks a visual perturbation was applied to the movement of the cursor consisting of a 45° clockwise rotation of the cursor trajectory with respect to its actual one. In each of these blocks subjects performed ten repetitions of three targets randomly presented on the screen. Subjects were divided in two groups (7 individuals each group). Group1 experienced targets NW, NE and E during the Ad1-Ad3 blocks and targets N, SE and SW during the Ad4-Ad6 blocks, while Group2 experienced the target-sets in the inverse order. After the Ad1-Ad6 blocks subjects performed four washout blocks each consisting of ten repetitions of three targets, this time without the visual perturbation. In the first two blocks of Washout (Wo1-Wo2) subjects reached to the targets trained in Ad1-Ad3, while in the second two blocks (Wo3-Wo4) they reached to the targets trained in Ad4-Ad6.

### Biomechanical analysis

The data recorded from the load cell was processed and analyzed to characterize the biomechanical patterns of adaptation and generalization of the two groups of subjects. Only the *x* and *y* components of the load cell were considered. The continuous load cell data recorded during each block were first low-pass filtered at 10 Hz using a 3^rd^ order Butterworth filter. After filtering, the load cell data of each reaching movement were segmented between the instant in which the cursor exited the central circle and the instant in which it reached the target circle. All segments were then time-scale normalized to 100 samples. We evaluated the adaptation and generalization patterns by calculating the Normal Curvilinearity (NC) of their reaching trajectories. NC was calculated for each length-normalized movement as the ratio between the length of each movement trajectory and the length of the straight line connecting the boundaries of the circles representing the center of the space and the target. A statistical analysis was performed to assess for differences in the average NC values calculated at the beginning of adaptation and generalization of all targets pooled together to assess for differences in NC due to generalization. This analysis was based on Mann-Whitney U test with significance level α set to 0.05.

### EMG signal recording and processing

EMG signals were recorded from the following 13 upper limb muscles: Brachiradialis (BRD), Biceps brachii short head (BSH), Biceps brachii long head (BLH), Triceps brachii lateral head (TLT), Triceps brachii long head (TLN), Deltoid Anterior (DANT), Medial (DMED) and Posterior (DPOST) heads, Pectoralis Major (PM), Inferior head of the Trapezius (TRAP), Teres Major (TMAJ) and Latissimus Dorsi (LD). EMG signals were recorded through a Delsys Trigno system (Delsys, US), sampled at 2000 Hz and synchronized with the load cell. All the subsequent analyses were performed in the Matlab environment (2014b, Mathworks, US). EMG signals were filtered in the 20Hz-400Hz band by using a 3^rd^ order digital Butterworth filter. Amplitude envelopes were then obtained by low pass filtering the full wave rectified EMGs with a 3^rd^ order Butterworth filter with a cut-off frequency of 10Hz.

Before muscle synergies extraction, all the calculated envelopes underwent an amplitude and time scale normalization procedure. Time scale was normalized by segmenting the EMG data based on the synchronized biomechanical data. Each center-out reaching movement was time-scale normalized by resampling the corresponding envelope on a fixed 100-samples time reference, representative of the percentage of execution of each reaching movement. The EMG envelopes of each subject were amplitude normalized by using the values extracted from the initial normalization block; during the normalization block, each participant reached three times to all the 8 targets, and the peak amplitude of each muscle during each movement was calculated. For each muscle the target yielding its maximal activation was identified. The reference normalization value for each muscle was then established as the average across the three peak values recorded across the repetitions of the target maximizing each muscle’s activity during the normalization block.

### Muscle synergies extraction at baseline

Muscle synergies were extracted from each participant at block B3. The NMF algorithm with multiplicative update rule^41^ was applied to the matrix D_13 × (40 × 100)_ of the EMG envelopes, containing the time scale and amplitude normalized EMGs from the 40 reaching movements (8 targets × 5 trials). For each subject, a synergy vectors matrix W_13xS_ and a matrix Ap_Sx(40 × 100)_ of the synergy activation patterns was obtained; a number of synergies from 1 to 13 was extracted, and S was defined as the lowest number of synergies able to explain an average (across subjects) fraction of variance higher than 90%.

### Synergies Activation Patterns reconstruction during motor adaptation using the fixed B3 synergies and statistical validation

For each subject, the EMG envelopes of all the subsequent adaptation blocks Ad1-Ad6 and washout blocks Wo1-Wo4 were reconstructed by using the set of baseline synergies extracted at B3 from the same subject. This was achieved by using a modified version of the NMF algorithm (namely NonNegative Reconstruction, NNR)^40^, able to reconstruct the matrix D by letting only the coefficients matrix AP update at every algorithm iteration, while keeping the W matrix fixed. The ability of the B3 synergies to account for the envelopes from all the trials was verified in two separate ways. First of all, the variance accounted for by the reconstruction (VAF_REC_) was quantified and statistically compared across trials. In second instance the obtained VAF_REC_ values were statistically compared to those expected from chance. For each subject and each block, 100 random synergy matrixes were obtained by shuffling the muscle components within each synergy, in order to obtain structure-less synergy matrixes W_SHUFFLE_. These matrixes were then used in NNR algorithm and obtain VAFSHUFFLE values from structure-less modules. We then tested for significant differences between VAF obtained from the reference synergies and those obtained from chance for each block. This analysis was based on Mann-Whitney U test with α = 0.05 and Bonferroni’s correction to take into account the repeated analysis on 11 blocks (from B3 to Wo4), for an effective α of 0.0045.

### Calculation of the preferred angles of the synergies APs

To evaluate the angular tuning of synergies APs, we calculated the preferred angle spanned by each synergy AP in the workspace at both baseline and at full adaptation for all 6 adapted/generalized targets. Preferred angles were calculated from the parameters of a cosine fit^28^ between the RMS of each synergy and the corresponding target position. Synergies APs were fitted using a linear regression in the form: *AP*(*θ*) = *β*_0_ + *β*_1_ cos(*θ*) + *β*_2_ sin(*θ*). The preferred angle of each synergy was then calculated from the fitting parameters as *ϑ* = *tan*^−^1(*β*_2_/*β*_1_). The preferred angles at baseline were calculated from the RMS of the average (across repetitions and subjects) APs at B3. Preferred angles at final adaptation were calculated by pooling together, for each group, the RMS of the average (across the last 5 repetitions of each target and across subjects) APs at the end of Ad3 and Ad6.

### Quantification of AP similarity between current and CW rotated targets

During the Ad1-Ad6 blocks, each synergy Ap_X_ towards a target X, as obtained from applying the NNR algorithm, was compared with the corresponding average Ap_X-45°_ towards the 45° CCW-rotated target at B3. Similarity was quantified by calculating the correlation coefficient between Ap_X_ and Ap_X-45°_ for each target. Specifically, the similarity was calculated between the average Ap_X-45°_ of the last five repetitions of a given target at B3 and the average AP calculated in either the first or last 5 repetitions of a movement in a certain block. To assess for differences in the final adaptation states (see **Results**, *Differential paths to adaptation*) the similarities were calculated between B3 and the end of Ad3 and Ad6. To assess for differences in generalization the similarities were calculated between B3 and the beginning of Ad1 and Ad4. In this latter analysis, Mann-Whitney U test was performed to statistically compare AP similarity between targets differently used as adaptation or generalization targets by the two groups. Linear correlation analysis was also used to assess for linear correlation between the similarities calculated between B3 and Ad1/Ad4 and the relative values of NC observed at Ad1/Ad4. Strength and significance of the linear correlation were assessed using Spearman’s coefficient. A statistical analysis was employed to assess for differences in the values of similarities between B3 and Ad1/Ad4 depending on the targets grouped by the difference in preferred angles between baseline and adaptation. This analysis was based on Kruskal-Wallis test with α = 0.05).

## Author contributions

G.S. conceived the study; G.S., C.D.M., J.D.S. and S.C. designed the study protocol; G.S., J.D.S. and M.Z. set up and executed the experiments; G.S., C.D.M. and J.D.S. analyzed the data; G.S. and C.D.M. drafted the manuscript; J.D.S., M.Z. and S.C. reviewed the draft.

## Competing interests

The authors declare that they have no competing interests.

## Data Availability

Please contact GS to obtain a copy of the data.

